# Using High Throughput DNA Sequencing to Evaluate the Accuracy of Serial Dilution Based Tests of Microbial Activities in Oil Pipelines

**DOI:** 10.1101/323139

**Authors:** K. Khanipov, G. Golovko, M. Rojas, M. Pimenova, L. Albayrak, S. Chumakov, R. Duarte, W.R. Widger, T. Pickthall, Y. Fofanov

## Abstract

Microbial activities have detrimental effects on industrial infrastructure. If not controlled, microbial presence can result in corrosion, biofilm formation, and product degradation. Serial dilution tests are routinely used for evaluating presence and abundance of microorganisms by diluting samples and culturing microbes in specific media designed to support microorganisms with particular properties, such as sulfate reduction.

A high-throughput sequencing approach was used to evaluate changes in microbial composition during four standard serial dilution tests. Analysis of 159 isolates revealed significant differences in the microbial compositions of sequential serial dilution titers and identified several cases where: (a) bacteria known to have a detrimental metabolic function (such as acid production) were lost in the serial dilution medium designed to test for this function; (b) bacteria virtually absent in the original sample became dominant in the serial dilution medium. These observations raise concerns regarding the accuracy and overall usefulness of serial dilution tests.

## 1. Introduction

Microbes are everywhere. Their presence can have detrimental effects on the longevity of industrial infrastructure (Joshi 2007). Microbial activities are associated with loss of product quality, biofilm formation, and microbiologically influenced corrosion (MIC). In 2013 the US spent over 452 billion dollars (over 2.7% of GDP) on corrosion related expenses of which at least 20% were associated with microbial activities (Koch et al. 2016; Koch et al. 2002). Standard approaches for protection against MIC include a combination of expensive alloys, protective coatings, mechanical cleaning, and treatment with biocides. The wrong choice of biocide, however, may lead to unexpected results. For example, using biocide ActiSEPT (sodium dichloroisocyanurate) in the presence of *Geotrichum* spp. (fungus) has been reported to significantly increase the microbiologically influenced corrosion of AISI 304 stainless steel instead of mitigating it (Stoica et al. 2012).

Several industry standards have been developed to identify the presence of microbes and their potential involvement in undesirable activities. The most frequently utilized standards: TM0106-2006 (NACE 2006), TM0194-2014 (NACE 2014) and TM0212-2012 (NACE 2012) were established by National Association of Corrosion Engineers (NACE International). The American Society for Testing and Materials (ASTM International) also introduced several standards to test microbial activity (e.g., D6469 and D6974). To date, these standards have been used to develop a variety of commercial testing kits such as fuel/oil MicrobMonitor2™ (ECHA Microbiology Ltd.), and series of biological activity reaction kits (BART™) for waterborne microbes (Drycon Bioconcepts Inc.). All these tests require the growth of microbes in specific media mimicking environmental conditions (e.g., aerobic or anaerobic). The quantity and capacity of microbes to perform specific undesired functions (e.g., acid production or sulfate reduction) are estimated by observing the outcomes of microbial activity across sequentially diluted samples. Due to the low initial abundance and relatively slow growth rate of some environmental microorganisms, serial dilution testing can take two to six weeks to complete.

Growing microbes in selective media for diagnostic purposes is also a standard practice in clinical microbiology. The goal of clinical diagnostics, however, is to confirm (detect) the presence of specific microbes of interest (usually pathogens), so over 600 specific growth media are routinely used for this purpose (Atlas and Snyder 2011; Jorgensen et al. 2015). In contrast, the serial dilution tests aim to detect the presence of microbes exhibiting specific functional properties, so the assumption behind this approach is that the most “important” microbes involved in detrimental activities (e.g., hydrogen sulfide or acid production) will survive and produce visible changes (such as color alteration) in the growth medium.

Serial dilution tests may result in the oversight of some important properties of microbial communities. Overwhelming evidence highlights the importance of synergistic relations between different microorganisms (bacteria, fungi, algae) in microbial communities (MC) (Beech and Sunner 2004; Videla 2001; Videla 2015). The nature of serial dilution tests, however, does not allow one to assess the effects of the collective behavior of microorganisms as only small portions of members of MC can survive in any given medium. As result, the microbiome’s biofilm formation abilities, which requires many different microbes to work together and usually a combination of aerobic (on the biofilm surface) and anaerobic (deep inside the biofilm) microorganisms cannot be detectable using serial dilution tests.

The purpose of the presented study was to use high-throughput sequencing technology to explore the accuracy of the four common serial dilution tests aiming to detect activities of general aerobic, aerobic and anaerobic acid producing, and sulfate reducing bacteria in oil pipes.

## 2. Materials and Methods

### 2.1. Sample collection

Sixteen samples were collected from the internal surfaces of two oil pipelines in the Southwestern United States. Fifteen samples were collected from a single pipeline at three locations several miles from each other. Samples were taken from the corroded pits at the six o’clock position of the pipeline and the adjacent non-corroded areas at the three, six, nine, and twelve o’clock positions. One additional sample was taken from the corrosion monitoring coupon of the second pipeline in the same geographical area (Table 1). Sample collection was performed with sterile swabs coated in anaerobic dilution solution (ADS) by Dixie Testing and Products, Inc. (Dixie 2013) and prepared according to NACE standard TM0194-2014 (NACE 2014) for field monitoring of bacterial growth. Specifically, swab samples were collected from 4 cm^2^ areas, placed into 10 ml tubes with ADS, and capped. The sample tubes were agitated to suspend the mixture with one milliliter of the solution and 9 ml of growth medium used for each of the four serial dilution tests. Sequential serial dilution titers were made by taking one milliliter of solution from the suspended mixture and transferring it into new dilutions. Samples were grown in eight titers of dilutions in the four media prepared according to NACE standard TM0194-2014 (NACE 2014): aerobic acid producing (Phenol Red Broth), anaerobic acid producing (Phenol Red Broth purged with nitrogen), general anaerobic (Thioglycolate), and sulfate reducing (API-RP38 modified). The lowest and highest dilution titers where microbial growth was visibly observed (solution changed color) can be found in Table S1 of the Supplementary Materials Document. All serial dilution samples were processed simultaneously.

**Table 1.** Sample locations and serial dilution (low/high) titers considered in the analysis.

| Sample ID | Pipeline | Location | Position in Pipe |
| --- | --- | --- | --- |
| 111 | 1 | 1 | Pit 6" |
| 112 |  |  | Away from Pit 6" |
| 113 |  |  | 12" |
| 114 |  |  | 3" |
| 115 |  |  | 9" |
| 116 |  | 2 | Pit 6" |
| 117 |  |  | Away from Pit 6" |
| 118 |  |  | 12" |
| 119 |  |  | 3" |
| 120 |  |  | 9" |
| 121 |  | 3 | Pit 6" |
| 122 |  |  | Away from Pit 6" |
| 123 |  |  | 12" |
| 124 |  |  | 3" |
| 125 |  |  | 9" |
| 211 | 2 | 4 | coupon |

### 2.2. High-Throughput DNA Sequencing

High-throughput sequencing of the V3-V4 16S ribosomal RNA gene region was performed using DNA isolates from each of 16 original samples, the corresponding anaerobic dilution solution (ADS), as well as isolates from serial dilution medias. To make sure that microorganisms in each sample are present in significant abundance, only media with microbial growth (the first and last titer) of each of the four dilution media were used in the analysis. DNA was isolated using the PowerSoil™ DNA isolation kit (Mo Bio Laboratories, CA). Sequencing libraries for each of the 159 isolates were generated using universal 16S rRNA V3-V4 region primers (Klindworth et al. 2013) in accordance with the Illumina 16S rRNA metagenomic sequencing library preparation protocol. Sequencing was done using an Illumina MiSeq instrument and resulted in 220,786 ± 6,721 DNA fragments (reads) per isolate with read lengths varying between 150 and 300 nucleotides.

### 2.3. Bioinformatics analysis

The analysis of the sequencing data was performed using the CLC Genomics Workbench 8.0.3 Microbial Genomics Module (http://www.clcbio.com). Sequencing reads containing unknown or low-quality nucleotides, library adapters, and chimeric reads have been excluded using the standard CLC Genomics Workbench sequencing data analysis protocol. The original sequencing reads for each sample were deposited in the sequence reads archive (SRA) at the National Center for Biotechnology Information (NCBI), project id: PRJNA302694. The filtered and trimmed reads are available by request.

To ensure consistency of the downstream analysis, the resulting high quality reads were uniformly trimmed to 135 bases. Reference based operational taxonomy units (OTU) assignment was performed using the SILVA SSU v119 97% database (Quast et al. 2013). Unassigned sequences identified in more than one copy across at least one dataset were placed into *de-novo* OTUs (with 97% similarity threshold) and aligned against the SILVA database with an 80% similarity threshold to assign higher level taxonomy associations. Detailed 16S rRNA gene-based profiles of each sample can be found in Table S2 of the Supplementary Materials Document. To address the concern that relatively short (135 nucleotides long) reads can be used to identify bacteria on genus taxonomy level, the identification of microorganisms has been performed on lowest sequence clusters (OTU) level. In this approach microorganism (regardless of their taxonomic identification) have been considered if the particular sequence has been present or absent in samples.

## 3. Results and Discussion

The taxonomic annotation of 159 sequenced isolates indicated the presence of 48 phyla, 183 classes, 407 orders, 766 families, 1,756 genera, 3,941 species, and 24,318 individual OTUs (most of which can be interpreted as distinct strains) of archaea and bacteria. The average numbers of operational taxonomic units (OTUs) for samples (α-diversity and entropy) (Figure 1 and Table S3 of the Supplementary Materials Document) reflect the presence of a very diverse microbial population in the original and ADS isolates. As expected, a significant loss of diversity was observed in serial dilution isolates due to the selective conditions of growth media. The lowest microbial diversity was observed in general anaerobic and aerobic acid producing media. Microbial communities in the sulfate reducing media appeared to be the most diverse out of all the dilution media (Figure 1), reflecting the less selective conditions for microbial survival in this media. The downstream comparisons of microbial compositions of individual isolates were focused on the factors able to affect the outcome of the serial dilution tests.

**Figure 1.**
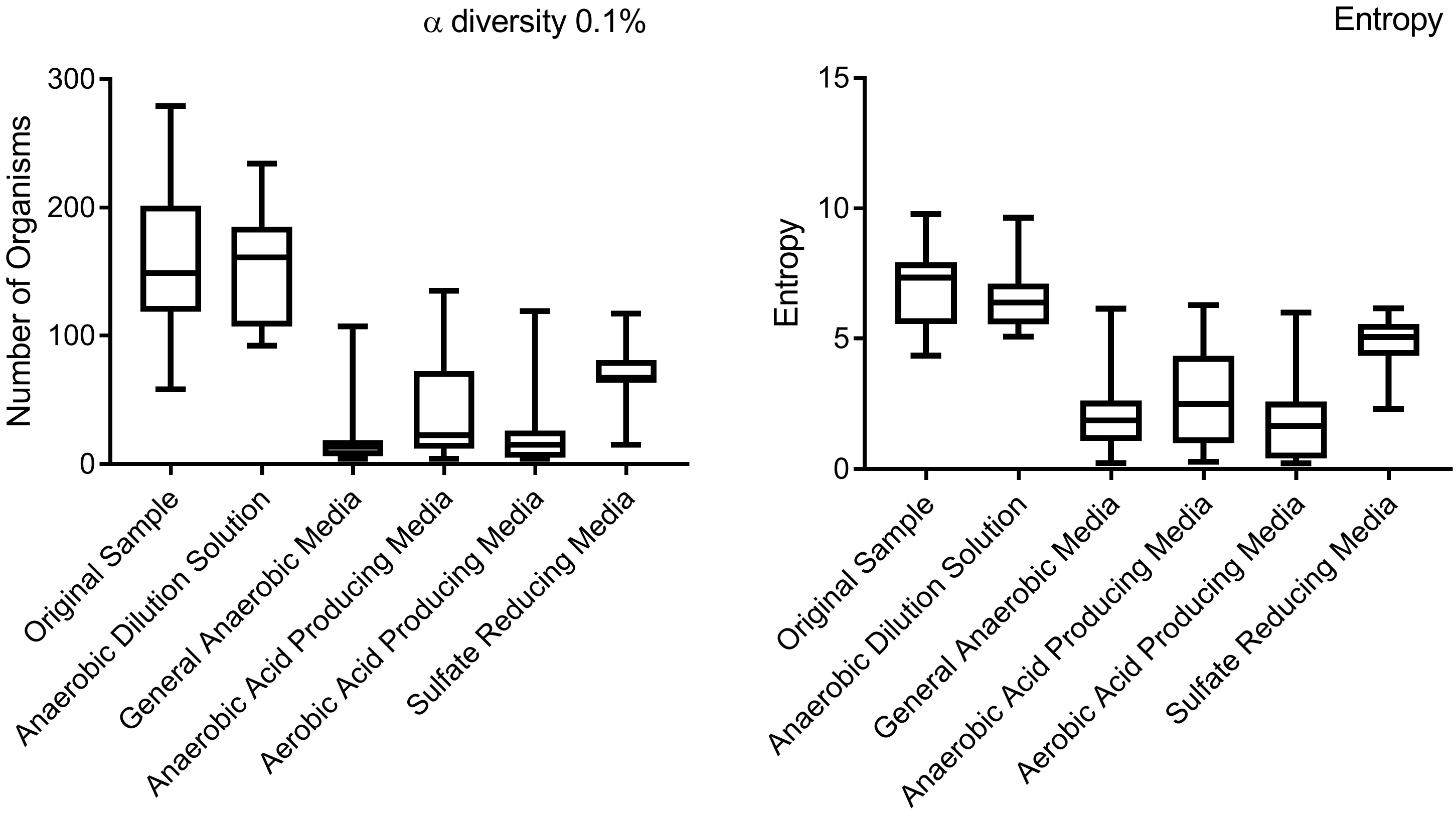
Changes in α diversity and entropy (Shannon diversity) of microbial compositions between original and serial dilution isolates.

### 3.1 Differences between original and anaerobic dilution solution samples

Preservation of original samples in the anaerobic dilution solution (ADS) is the first step of every serial dilution test. Some microorganisms (e.g., obligate anaerobes which cannot tolerate the presence of oxygen), however, may not survive the stress of the sample handling, transportation, and ADS preparation. The presence of favorable conditions in ADS for some microorganisms can also alter the microbial abundance in the sample: under favorable conditions, some bacteria can double their biomass every 30 minutes. It is also possible that superfluous microorganisms may appear in the sample due to contamination. The basic assumption of the serial dilution tests, however, is that all these effects introduce only minor bias into the microbial composition carefully enough can represent the microbial composition of the original sample.

Comparison of the DNA profiles of corresponding original and ADS isolated revealed significant differences in microbial composition. Low values of the Pearson correlation coefficient: 0.652 ± 0.057 on OTU and 0.63 ± 0.064 on species levels (Figure 2 and Table S4 of the Supplementary Materials Document) suggested that substantial changes in microbial abundance occur in the microbial population during the ADS preparation step, potentially negating the ability of the microbial community to exhibit functionality during serial dilution testing.

**Figure 2.**
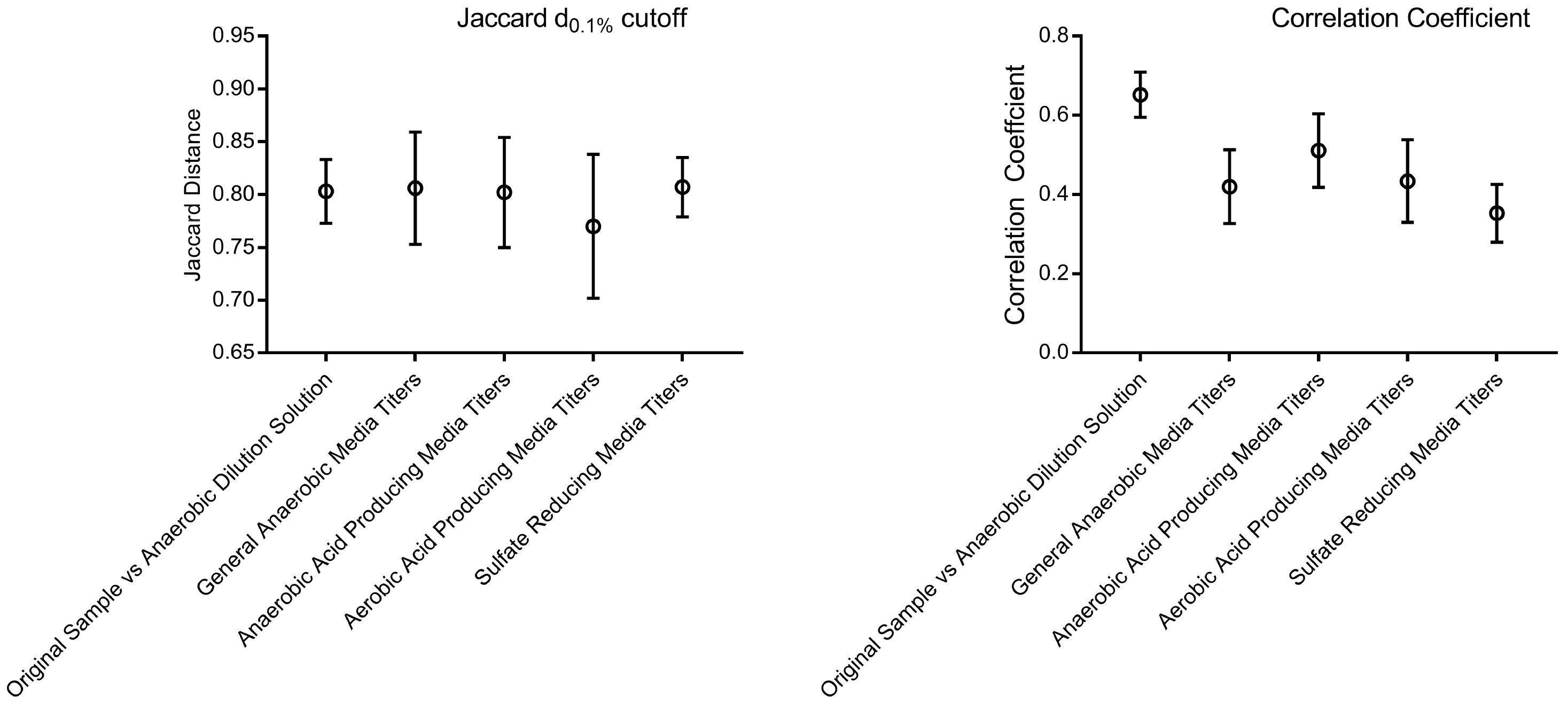
The low similarity between microbial profiles in original samples and ADS as well as between corresponding titers of serial dilution medias.

Since some microorganisms can grow rapidly, one may argue that as long as microorganisms are still present above certain minimal abundance levels, the changes in its relative abundance between the original and ADS samples may not affect this organism’s ability to perform its metabolic role in the microbial community. For example, to fulfill its role, nitrogen fixing bacteria can be present in as low as 0.1% of relative abundance (Saito et al. 2011). To include all microorganisms which may contribute to the metabolic profile of the microbial community, the analysis of the bacterial presence/absence profile was performed using three different relative abundance thresholds (1%, 0.1%, and 0.01%). Comparison of the bacterial presence/absence profiles of the original and ADS samples at the OTU level revealed that up to 80% of microorganisms can be “lost” or “gained” (appeared above or below the relative abundance threshold) during the ADS preparation step. The similarity score between bacterial presence/absence profiles of corresponding original and ADS samples calculated using Jaccard distance (Levandowsky and Winter 1971) (Figure 2 and Table S4 of the Supplementary Materials Document) demonstrates that loss and gain of bacteria can be observed on species, genus, and family taxonomic levels.

Considering a strict asymmetrical dual abundance threshold (over 1% for presence and below 0.001% for absence), gain and loss of bacteria between the corresponding original and ADS were detected for 53 organisms (OTU level) in nine out of sixteen samples (Table S5 of the Supplementary Materials Document). It is important to mention that loss of some organisms, such as *Simplicispira sp.* (acid producing facultative anaerobes), can be interpreted as the inability of the ADS to support the survival of some acid-producing organisms.

### 3.2 Differences between microbial composition in first and last serial dilution titers

The main purpose of the use of serial dilutions in MIC detection is to quantify the presence of harmful microbes in the samples. The underlying assumption of the dilution experiments is that the same microorganism(s) relevant for the purpose of the test will grow in all the titers, so the different levels of titration (dilution) of the original sample can elucidate the initial concentration of these microorganism(s). Comparison of the bacterial DNA profiles of the corresponding first and last titers of each of the four types of serial dilution tests suggests low stability of the microbial community between titers. Both, the Pearson correlation coefficients and Jaccard distances suggested significant dissimilarity between corresponding titers (Figure 2 and Table S6 of the Supplementary Materials Document). Particularly low correlation between titers (*p*=0.35) were observed for the sulfate reducing bacteria medium at the OTU level. In thirteen out of the sixteen samples, at least one of the OTUs virtually “disappeared” between the first and last titers (relative abundance was detected above 30% in the first and dropped below 1% in the last titer).

A “typical” example of how bacterial compositions between the first and the last dilutions of sulfate reducing bacteria tests have changed on the taxonomic rank of *order* is presented in Figure 3 (complete sample-by-sample taxonomy profiles can be found in the Supplementary Materials Document). As one can see, bacteria in the first dilution titer were distributed between the dozens of taxa. The two most abundant taxa: *Clostridiales* and *Enterobacteriales*, were present in relative abundance of 16.26% and 12.85%, respectively. The last titer however, showed the absolute dominance of *Bacillales* (71.39%), of which about 35% was contributed by *Bacillus weihenstephanensis* spp. (identified on species level due to its distinctiveness from other *Bacillales* 16S sequence), whose abundance in the first titer of the dilution was only 0.022%. This observation suggested that this bacterium was not part of the original microbial community but appeared there by accident (e.g., contamination). Another example of changes in bacterial abundance includes *Pseudomonadales* whose relative abundance increased from 2.5% to 7.5% and *Rhodospirillales* whose abundance increased from 0.04% to 1.12%.

**Figure 3.**
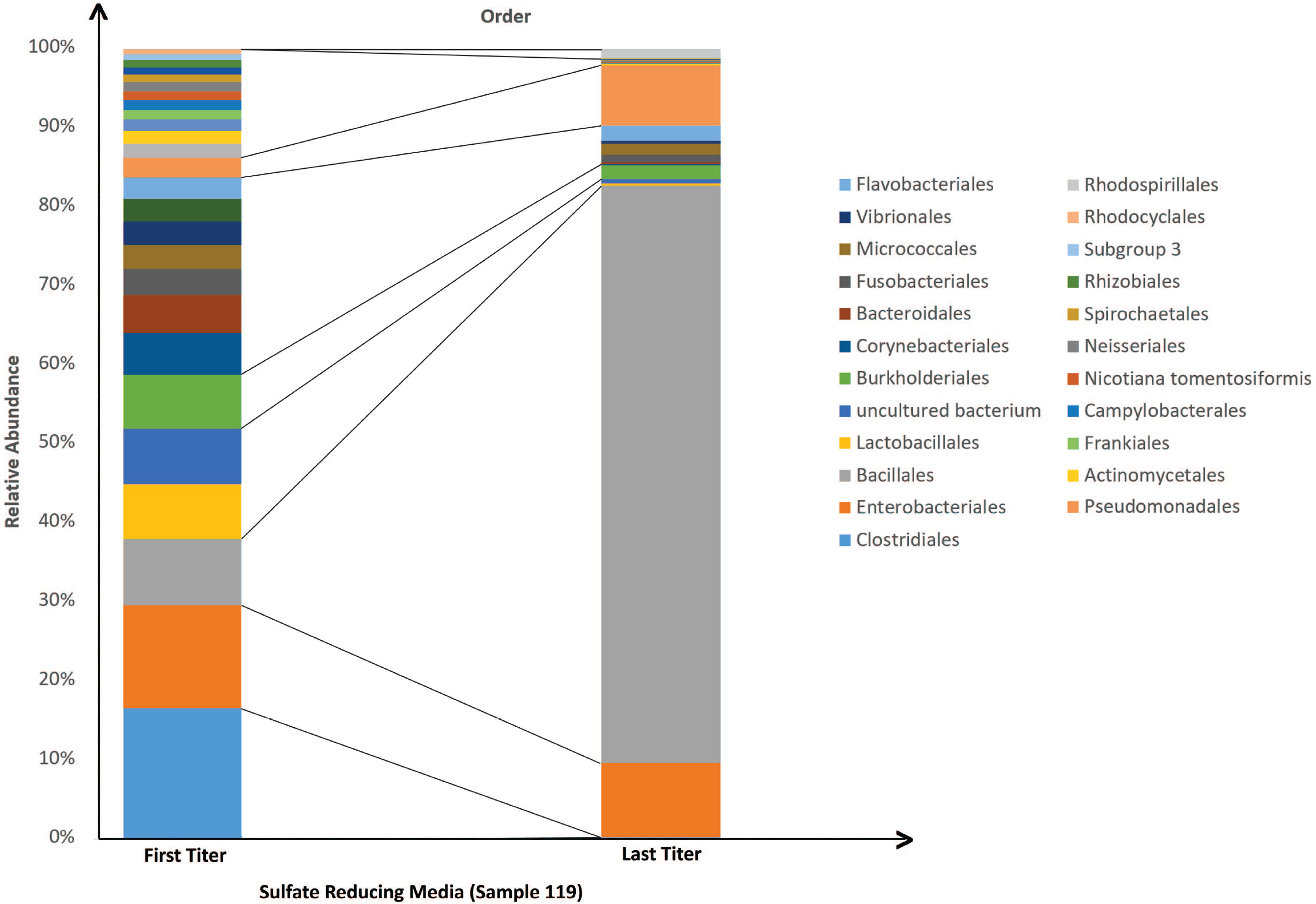
Changes in microbial composition (taxonomical level of order) between first and second sulfate reducing serial dilution titer isolates (sample 119).

The overall comparison of microbial compositions between the first and the last titers of the same types of serial dilutions suggested that the stability of the microbial communities across titers in the serial dilution tests is very low.

### 3.3 False positive results: bacteria absent in the original sample and detected in the serial dilution test

Perhaps the most misleading outcome of the serial dilution test is when harmful microorganisms absent in the original sample are reported as a result of the test (false positive result). In every analyzed sample, at least one OTU that was virtually non-existent (present in relative abundance <0.001%) in the original sample was detected in the serial dilutions isolates in significant (above 5%) abundances (Tables S7a-S7d and Table S8 of the Supplementary Materials Document). In 68% of the dilution samples (up to 25% anaerobic acid producing, 63% general anaerobic, 25% aerobic acid producing, and 68% sulfate reducing bacterial media) at least one organism (OTU) was identified to be a false positive. In some cases, such OTUs even dominated (above 50% relative abundance) the sample. While the origin of the false positive organism can be both contamination and growth from extremely low abundance (and for this reason not contributing to the metabolic activity of the original microbial community) the fact that different organisms grow in functionally relevant abundances in all the serial dilution media in all sixteen samples suggests that this is not a random and statistically insignificant incident. Since all samples have been collected, preserved, and processed simultaneously, so contamination (if it occurred) must introduce the same combination of external microorganisms in all the samples. The fact that different “false positive” microorganisms have been detected in same types of media (but different samples) suggests that these organisms are originating from the samples.

### 3.4 False negative results: bacteria of interest present in the original sample and undetected in the serial dilution test

A false negative outcome of a test is when the microorganisms capable of performing the evaluated function (such as acid production) are present in the original sample but do not grow in the serial dilution media. The analyses performed have identified a number of cases where bacteria with metabolic activities known to be associated with the progression of MIC were completely lost in serial dilution media (Table 2). This includes *Bacillus funiculus,* an iron and sulfur reducing bacteria (Ajithkumar et al. 2002). At least one original sample was found to contain *Rhodococcus tukisamuensis*, an acid-producing bacteria (Matsuyama et al. 2003) which was lost in all corresponding dilution titers. Likewise, *Shewanella sp.* and *Aeromonas sp.,* synergetic iron reducers that require each other presence to grow (Knight et al. 1996), were found only in the original samples. Table S9 of the Supplementary Materials Document contains a large list of organisms capable of performing corrosion-related metabolic activities that were present in original samples but not identified in any of the serial dilution titers.

**Table 2.**
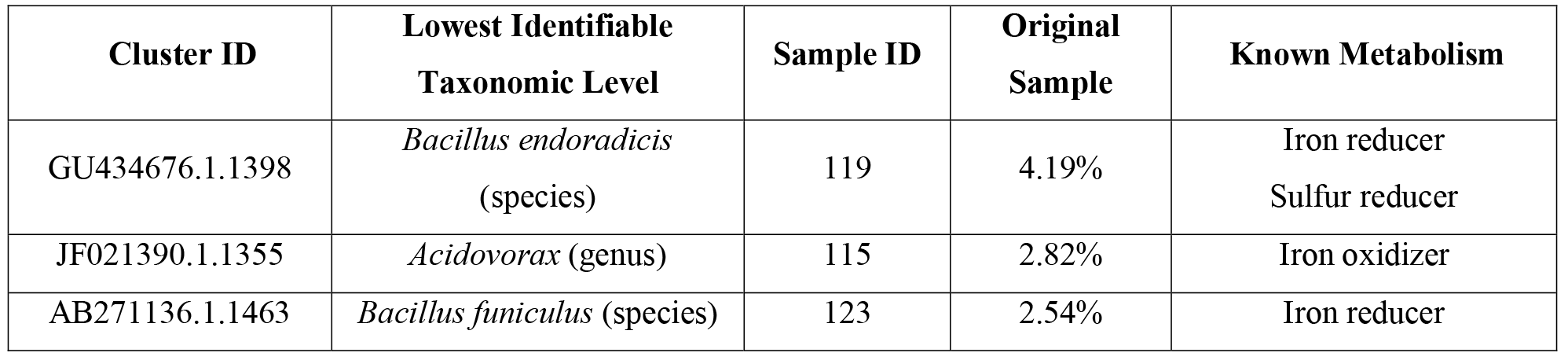
Bacteria with corrosion associated metabolic pathways lost in serial dilutions.

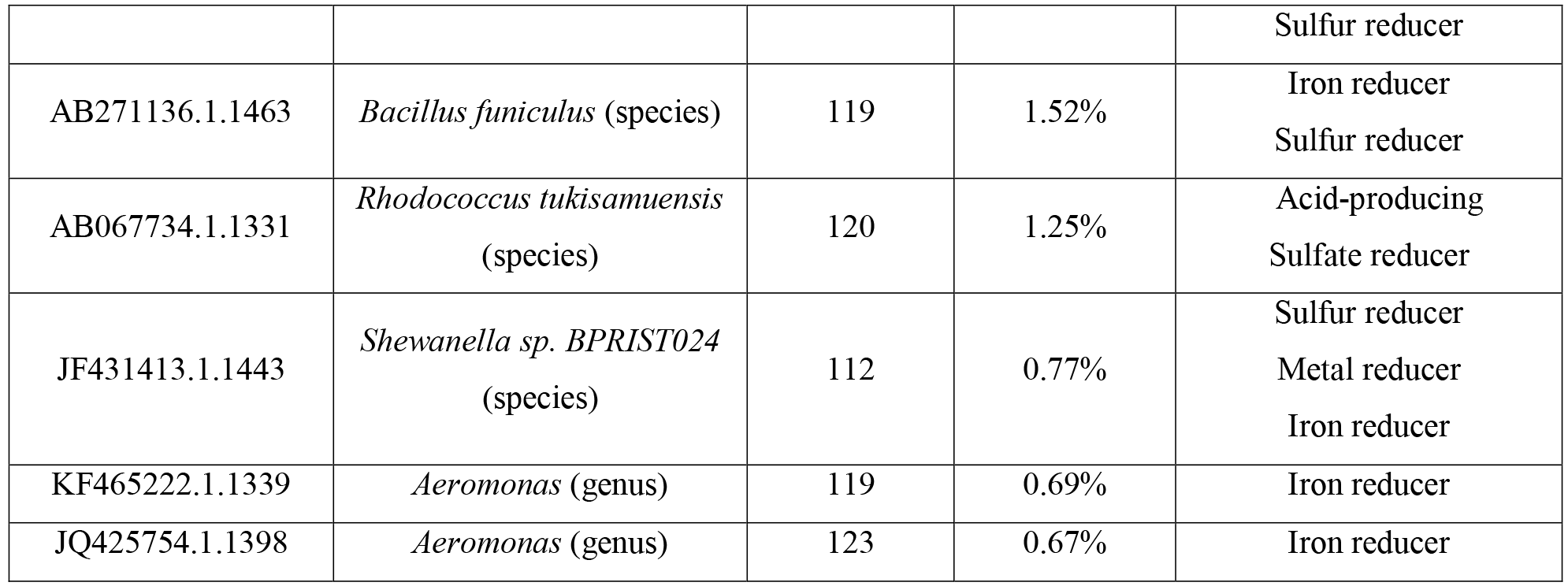

### 3.5. Potential biases of the HTS-based metabarcoding approach

Many external factors, from inappropriate sample handling (such as exposure of samples to air) to dilution solution preparation, can affect the outcome of both serial dilution and HTS based tests. The advantage of the HTS approach is that in contrast with serial dilution it can detect the presence of dead bacteria which did not survive the sample collection and processing steps. It is necessary to keep in mind that, regardless of recent progress in biomedical applications, HTS technology is still in its infancy in industrial applications. Amplification of selected genes such as 16S rRNA (for bacteria), ITS (for fungi), and COI (for microscopic eukaryotes), sometimes referred to as “metabarcoding”, (Loeffelholz and Fofanov 2015) has its limitations in the assessment of the metabolic or functional profile of the microorganisms: the presence/absence profile of many genes associated with specific metabolic activities may vary between individual strains of the same species. The complete functional profile of microbial community can be done using whole genome shotgun sequencing of total DNA and RNA isolated from the sample but this approach is still expensive and data analysis is limited by the absence of comprehensive reference databases. Another challenge for HTS technology is the presence of exoDNA and DNA from dead bacteria which can introduce bias into the results of the DNA (but not RNA) analysis, if samples are taken after recent biocide treatment. These challenges, however, are expected to be resolved in the near future.

## 4. Conclusion

The comparison of microbial compositions between original samples, anaerobic dilution solutions, and corresponding titers of serial dilutions revealed significant concerns regarding the overall accuracy of the serial dilution testing approach and its ability to evaluate undesired microbial activities. The analyses performed suggested that bacteria involved in undesired activities can not only change their relative abundance but can also be completely lost (false negative) or acquired (false positive) during the serial dilution tests. Presented data also contradicts the underlying assumption of the serial dilution experiments that the same microorganism(s) relevant for the purpose of the test will grow proportionally in all the titers. While the analysis was performed using samples collected from oil pipelines, similar results can be expected for other types of samples (water, fuel, and corrosive biofilms of various types of alloys). The latest progress in high-throughput sequencing technology, fast turnaround (2-3 days), and comparable costs to alternative testing methods (e.g. serial dilution tests) makes the DNA sequencing approach a substantial complement, and potential future alternative, to the existing standard testing practices.

## Conflict of Interest

The authors declare that they have no competing interests.

## Ethical approval

This article does not contain any studies with human participants or animals performed by any of the authors.

## Funding

YF, GG, MR, KK, and LA work was partially supported by The Sealy Center for Structural Biology & Molecular Biophysics and the Institute for Human Infections and Immunity (IHII) at the University of Texas Medical Branch (UTMB) and Texas Advanced Computing Center (TACC). SC work was supported in part by CONACYT (Mexico) project No. 84358

